# Resistance to *Candidatus* Liberibacter solanacearum in tomato landraces from Mexico

**DOI:** 10.1101/605188

**Authors:** José A. Garzón-Tiznado, Carlos A. López-Orona, Luciano Castro-Espinoza, Sixto Velarde-Félix, Marely G. Figueroa-Pérez, Jesús E. Retes-Manjarrez

## Abstract

*Candidatus* Liberibacter solanacearum (*CLso*) is an economically important plant-pathogen of tomato (*Solanum lycopersicum*) crops in the United States, Mexico, Central America, and New Zealand. Currently, there are no reports of resistance to *CLso* in tomato cultivars. Identification and development of *CLso*-resistant cultivars may offer the most efficient way to manage this tomato disease. Resistance of 46 tomato landraces collected in different regions of Mexico, representing a wide range of genetic variability from this country was evaluated. Two assays were done in consecutively years to assess the resistance to *CLso* under greenhouse conditions. Plants from both tests were inoculated with *CLso* through 20 *Bactericera cockerelli* insects per plant. In the first trial, landraces FC22 and FC44 showed a significantly higher proportion of resistant plants, less symptoms severity, and longer incubation time, followed by landraces FC40 and FC33 compared with the rest of the 42 landraces and 2 susceptible cultivars 60 days post inoculation (dpi). In the second assay, only landraces FC22 and FC44 had again significantly higher proportion of resistant plants, less symptoms severity, relative lower *CLso* titers, and longer incubation time in comparison with landraces FC40 and FC33 and the two susceptible cultivars 60 dpi, corroborating their resistance to *CLso*. Presence of *CLso* DNA in all resistant plants from both assays discards scape plants and indicates that the methodology used was adequate to discriminate between resistant and susceptible plants. These results confirm that landraces FC22 and FC44 are promising resistant sources for the development of *CLso*-resistant cultivars of tomato.

**Author summary:** The bacterium “*Candidatus* Liberibacter solanacearum” (*CLso*) is an important plant-pathogen of tomato crops in the United States, Mexico, Central America, and New Zealand. Tomato growers are lacking of cultivars with resistance to this pathogen and the development of resistant cultivars of this crop would make a sustainable business for these growers and healthy tomato consumption for humans. Tomato landraces from countries that are center of domestication of cultivated crops like Mexico, are potentially sources of resistance to plant-pathogens. Therefore, two tests were done looking for resistance sources to this pathogen and we found two tomato landraces (FC22 and FC44) showing high level of resistance to *CLso* because they had significantly higher resistant plants, less symptoms severity, lower *CLso* DNA concentration, and delay of the first symptoms in the inoculated plants in comparison with the two commercial cultivars and 44 tomato landraces collected from Mexico 60 days post infection. These landraces are promising resistant sources for the development of *CLso*-resistant cultivars of tomato.

## Introduction

*Candidatus* Liberibacter solanacearum (*CLso*) is an alphaproteobacterium associated with “tomato permanent yellowing” and “tomato decline” [1-4], a disease causing economic losses to the tomato crop sector in Oceania, Central and North America [1, 4-8]. The tomato/potato psyllid *Bactericera cockerelli* (Šulc) is responsible for transmission of the bacterium in the field. For this reason, the bacterial name “*Candidatus* Liberibacter psyllaurous” was also a proposed species name [9]. CLso has different solanaceous hosts like *Solanum lycopersicum* L., *Solanum tuberosum* L., *Solanum melongenea* L., *Capsicum annuum* L., *Nicotiana* spp L., *Physalis* spp. L., *Solanum elaeagnifolium* L., and apiaceous like *Daucus carota, Apium graveolens*, and *Pastinaca sativa* [1,2,10,11].

In the last 15 years, *CLso*, which was first described from leaves of *Solanum tuberosum* plants in New Zeland [12], has become more important worldwide for its high aggressiveness, increasing geographical distribution, wide host range, and because no commercial cultivars of potato and tomato resistant to this bacterium have been reported so far [1,8,10,11,13]. In North and Central America, the main symptoms of this disease in tomato plants are the following: overall chlorosis, severe stunting, leaf epinasty, leaf filimorphism or elongated leaves, leaf rolling, crispy leaf, purple discoloration of veins, excessive branching of axillary shoots, flower abortion, and deformation of the fruits [1-4, 14].

The management of this bacterium has been based mainly on the chemical control through the use of insecticides against the vector insect. This method has been partially effective, costly, and represents a biohazard [15,16]. Furthermore, this method can also contribute to the development of resistance in vector populations [17]. An effective alternative, without bio-risk, and accepted for the integrated management of *CLso* is the development of genotypes resistant to this group of pathogens [13,16,18,19]. In tomato crops, control of other diseases, like TYLCV, transmitted by vectors, through resistant cultivars has been widely used successfully around the world [21].

The first step for the development of disease-resistant cultivars is the screening of wild and/or domesticated genetic resources, to be used afterwards in the genetic breeding programs of agricultural crops [20]. The second desirable step is to analyze the genetic base and heritability of the target trait to design the best breeding model for the introgression of the desirable trait into cultivated background, and the third step would be to carry out the plan [21].

Mexico is considered a domestication center for tomato [22-24]. Several studies indicate that *Lycopersicon esculentum* var. *cerasiforme* Dunal and other tomato landraces are distributed along the Mexican country [25-27]. Nevertheless, there is scarce information regarding the genetic potential of these tomato relatives and its direct usefulness in tomato breeding programs [27].

Wild relatives and landraces of *Solanum lycopersicum* have been reported as sources of resistance to pests and diseases [28,29]. The wide range of variability that has been reported for these genetic resources worldwide can be used to solve different problems such as resistant genes to diseases [30-32]. The wild relatives and landraces of tomato are a valuable genetic resource and have been successfully used as resistance sources against *CLso*; for example, [14] and [33] found resistance to *CLso* in different accessions of *S. lycopersicum* from Mexico. However, no tomato cultivars resistant to this bacterium have been described in Oceania, Central and North America so far. This could be due to the lack of new sources of resistance and/or studies on the genetic basis of the resistant trait to *CLso*. Therefore, it is important to continue looking for new sources of resistance that support breeding programs in the development of tomato resistant cultivars to this important pathogen.

The objective of the present study was to identify resistance sources against *CLso* using landraces of *S. lycopersicum* from different regions of Mexico, to contribute to the development of tomato cultivars resistant to *CLso*.

## Results

### *CLso* identification

The samples taken from the inoculum source of tomato plants were positive for the amplification of the predicted 1168 bp fragment corresponding to the *CLso* bacterium (Fig 1). The sequence of the amplified fragment showed 100% nucleotide identity with the KF776420 isolate of GenBank corresponding to a sample of *CLso*-infected *B. cockerelli* from Guanajuato [39] and 99% nucleotide identity with isolates KF776422, EU918197 and FJ939136 tomato plants infected with this bacterium from the state of Sinaloa, and Texas (USA), respectively. The phylogenetic analysis of the *CLso* isolated sequence from the inoculum source grouped the strain used in this study within the *CLso* species, and clearly differentiated it from other species of the genus *Candidatus* Liberibacter, as well as from their respective vectors (Fig 2).

**Fig 1.**
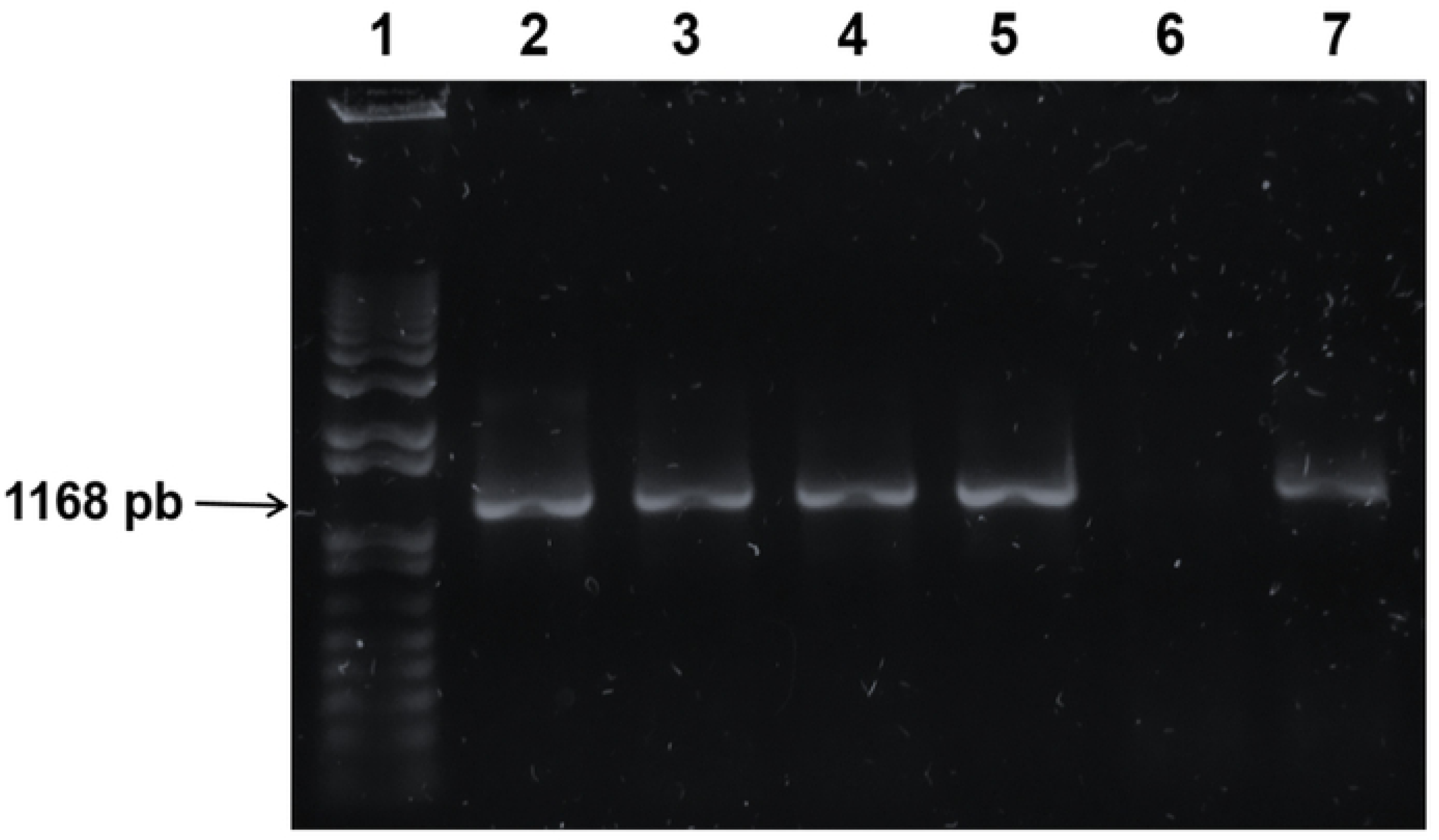
PCR products obtained from genomic DNA of *B. cockerelli* insects and tomato plants inoculated with *CLso* with the primer pair Oa2/OI2CF and Oa2/OI2CD, viewed on a 1.0% agarose gel. Lane-1 “Hyperladder 1 kb Plus”; lanes 2-3: DNA samples of *B. cockerelli*; lanes 4-5: DNA extracted from tomato CLso-infected plants; lane 6: DNA sample of non-inoculated tomato plant (negative control); lane 7: DNA sample from the inoculum source (positive control).

**Fig 2.**
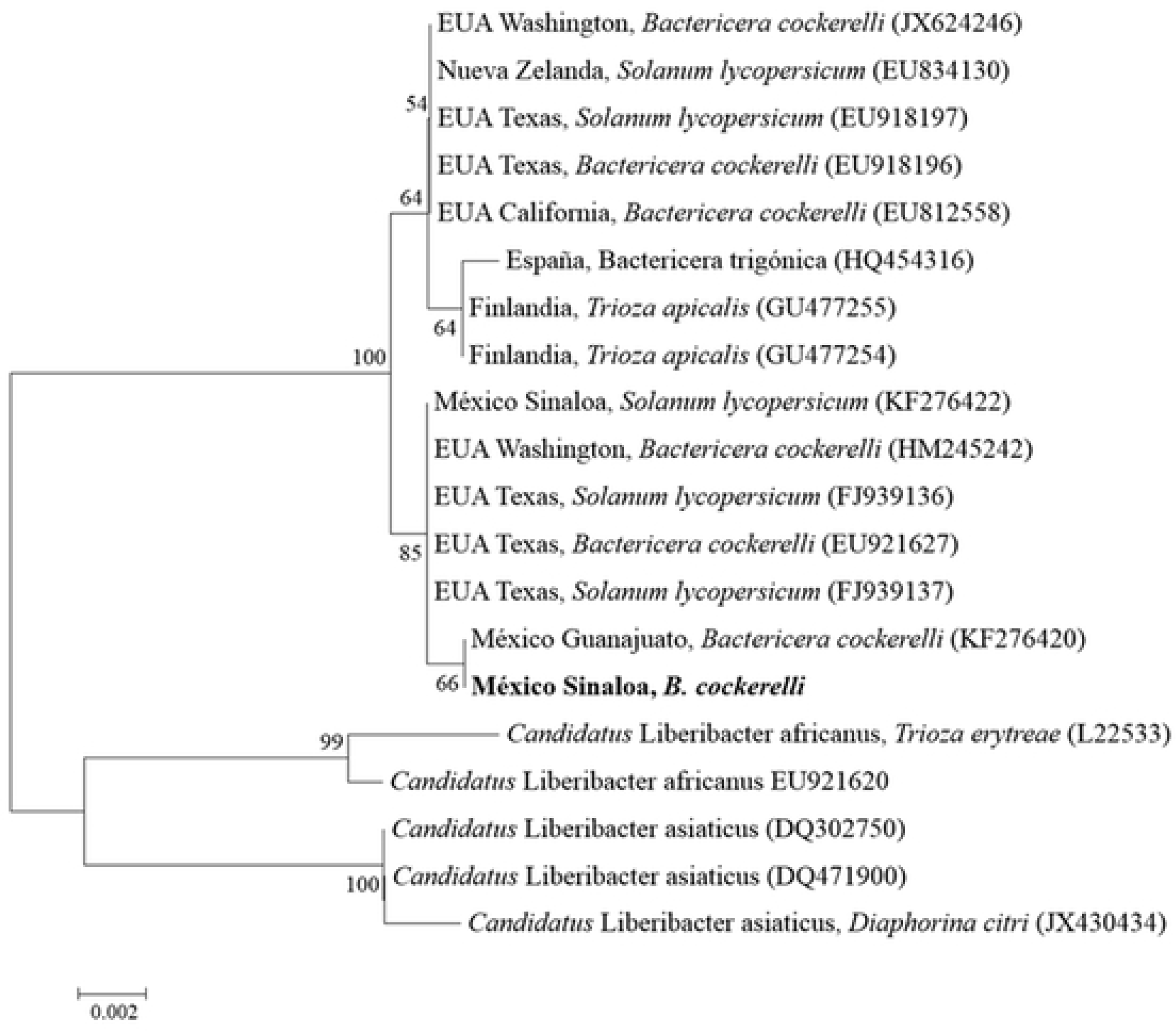
Phylogenetic dendogram using neighbor-joining method based on the alignment of partial nucleotide sequences of the 16S gene of *Candidatus* Liberibacter solanacearum and that of selected *CLso* strains (Mexico Sinaloa, *B. cockerelli*) included in both resistance assays. Values at the nodes represent the percentage bootstrap scores (1000 replicates), showing only those values above 50.

**Fig 3.**
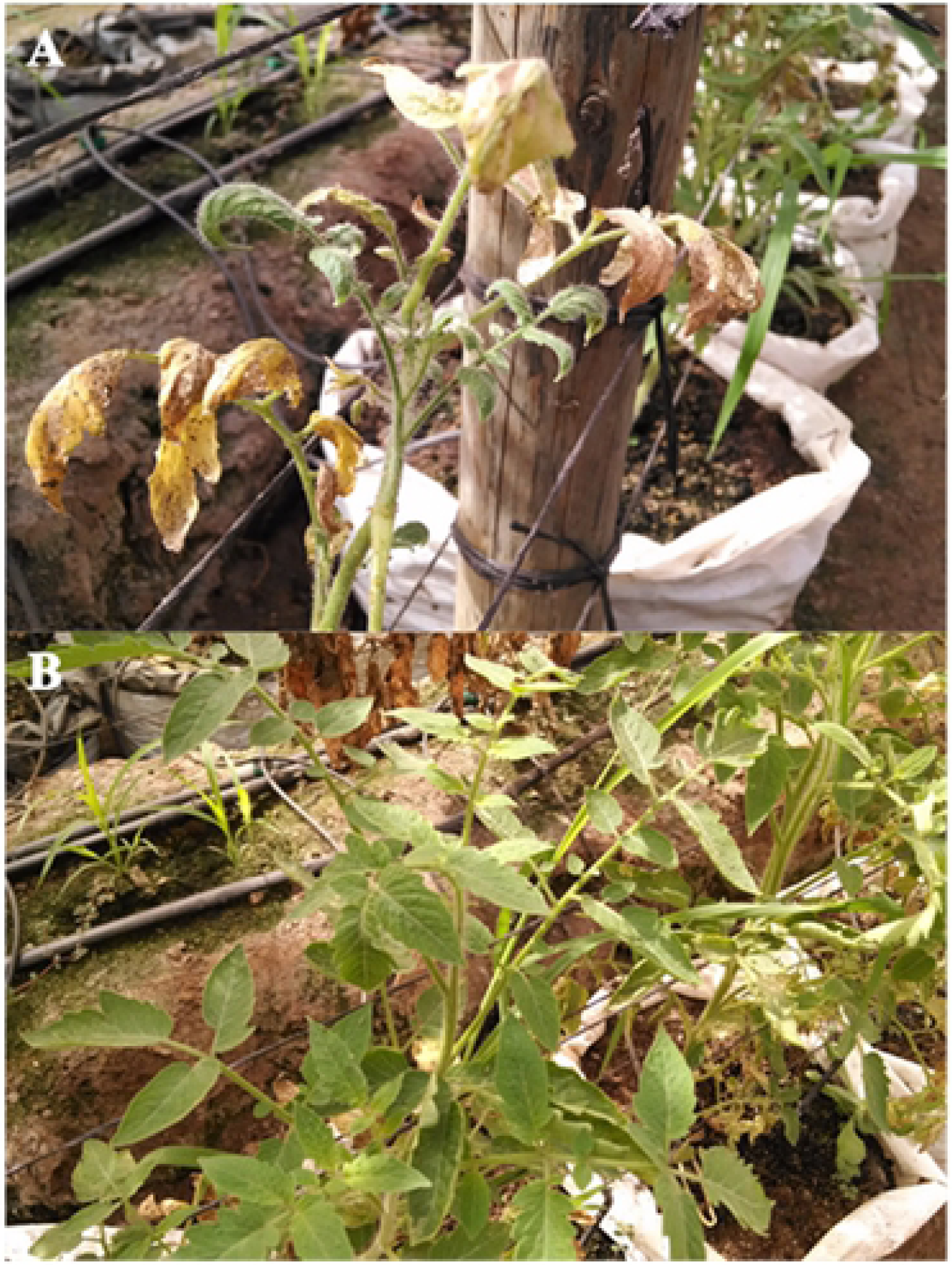
Susceptible tomato plant to *CLso* of the cultivar Pony Express (A), and resistant plant to *CLso* of the FC22 landrace (B). Pictures were taken 60 days post inoculation.

### First resistance assay

Most of the inoculated landraces were susceptible after 60 days post inoculation (dpi), although the percent of resistant plants, *CLso* symptoms severity, and longer incubation time varied significantly for all these parameters among genotypes in this assay (*P* < 0.0001) “Table 1”. Landraces FC22 and FC44 showed a significantly greater percent of resistant plants, lowest level of *CLso* damage, followed by the populations FC40 and FC33 with a percentage of resistant plants of 100, 100, 25, and 20 and an average of symptoms of 1.3, 1.9, 6.2, and 6.6, respectively. The rest of the landraces and susceptible Pony Express and Reserva controls were highly susceptible to *CLso* “Table 1”.

**Table 1.** First resistance assay of 44 tomato landraces and the cultivars Pony Express and Reserva inoculated with *CLso* through *B. cockerelli*.

| Origin | Genotype | NPR/TPP | Mean | Incubation | Range |
| --- | --- | --- | --- | --- | --- |
| Michoacan | FC26 | 20/20 (0%) | 8.9 A | 21.5 D | 7-9 |
| Jalisco | FC25 | 20/20 (0%) | 8.9 A | 21.5 D | 7-9 |
| Nayarit | FC13 | 20/20 (0%) | 8.9 A | 21.4 D | 7-9 |
| Michoacan | FC27 | 20/20 (0%) | 8.8 AB | 21.5 D | 7-9 |
| Jalisco | FC18 | 20/20 (0%) | 8.8 AB | 21.4 D | 7-9 |
| Jalisco | FC19 | 20/20 (0%) | 8.8 AB | 21.4 D | 7-9 |
| NA | Reserva | 20/20 (0%) | 8.7 ABC | 22.3 D | 7-9 |
| Nayarit | FC15 | 20/20 (0%) | 8.7 ABC | 21.4 D | 5-9 |
| Jalisco | FC24 | 20/20 (0%) | 8.7 ABC | 21.5 D | 7-9 |
| Nayarit | FC10 | 20/20 (0%) | 8.7 ABC | 22.1 D | 7-9 |
| Guerrero | FC38 | 20/20 (0%) | 8.6 ABC | 21.5 D | 7-9 |
| Sinaloa | FC02 | 20/20 (0%) | 8.6 ABC | 21.4 D | 5-9 |
| Sinaloa | FC09 | 20/20 (0%) | 8.6 ABC | 22.6 D | 7-9 |
| Michoacan | FC28 | 20/20 (0%) | 8.6 ABC | 21.5 D | 7-9 |
| Nayarit | FC12 | 20/20 (0%) | 8.6 ABC | 22.9 D | 7-9 |
| Michoacan | FC31 | 20/20 (0%) | 8.6 ABC | 21.5 D | 5-9 |
| Sinaloa | FC01 | 20/20 (0%) | 8.6 ABC | 21.8 D | 7-9 |
| Michoacan | FC29 | 20/20 (0%) | 8.5 ABC | 21.5 D | 7-9 |
| Michoacan | FC32 | 20/20 (0%) | 8.5 ABC | 21.5 D | 7-9 |
| Jalisco | FC20 | 20/20 (0%) | 8.5 ABC | 21.5 D | 7-9 |
| Michoacan | FC30 | 20/20 (0%) | 8.5 ABC | 21.5 D | 5-9 |
| Guerrero | FC35 | 20/20 (0%) | 8.4 ABC | 21.5 D | 7-9 |
| Guerrero | FC34 | 20/20 (0%) | 8.3 ABCD | 21.5 D | 7-9 |
| Sinaloa | FC03 | 20/20 (0%) | 8.3 ABCD | 22.0 D | 7-9 |
| Oaxaca | FC43 | 20/20 (0%) | 8.3 ABCD | 21.5 D | 7-9 |
| Oaxaca | FC41 | 20/20 (0%) | 8.2 ABCDE | 21.5 D | 7-9 |
| Nayarit | FC11 | 20/20 (0%) | 8.2 ABCDE | 22.4 D | 7-9 |
| Sinaloa | FC06 | 20/20 (0%) | 8.2 ABCDE | 22.3 D | 7-9 |
| Sinaloa | FC04 | 20/20 (0%) | 8.2 ABCDE | 22.1 D | 5-9 |
| Chiapas | FC45 | 20/20 (0%) | 8.2 ABCDE | 21.9 D | 7-9 |
| Nayarit | FC16 | 20/20 (0%) | 8.2 ABCDE | 22.5 D | 7-9 |
| Sinaloa | FC08 | 20/20 (0%) | 8.0 ABCDEF | 21.5 D | 7-9 |
| NA | Pony Express | 20/20 (0%) | 8.0 ABCDEF | 21.7 D | 7-9 |
| Oaxaca | FC39 | 20/20 (0%) | 7.8 BCDEFG | 21.5 D | 5-9 |
| Chiapas | FC46 | 20/20 (0%) | 7.7 CDEFGH | 21.5 D | 5-9 |
| Sinaloa | FC07 | 20/20 (0%) | 7.7 CDEFGH | 24.2 D | 5-9 |
| Nayarit | FC17 | 20/20 (0%) | 7.3 DEFGHI | 23.7 D | 5-9 |
| Guerrero | FC36 | 20/20 (0%) | 7.2 EFGHIJ | 21.5 D | 5-9 |
| Guerrero | FC37 | 20/20 (0%) | 7.0 FGHIJ | 21.5 D | 5-9 |
| Oaxaca | FC42 | 20/20 (0%) | 6.9 GHIJ | 21.5 D | 5-9 |
| Sinaloa | FC05 | 20/20 (0%) | 6.8 GHIJ | 23.6 D | 5-9 |
| Jalisco | FC23 | 20/20 (0%) | 6.8 GHIJ | 21.5 D | 5-9 |
| Nayarit | FC14 | 20/20 (0%) | 6.7 HIJ | 21.5 D | 5-9 |
| Jalisco | FC21 | 20/20 (0%) | 6.3 IJ | 21.5 D | 5-9 |
| Guerrero | FC33 | 16/20 (20%) | 6.6 IJ | 25.6 C | 3-9 |
| Oaxaca | FC40 | 15/20 (25%) | 6.2 J | 27.4 C | 3-9 |
| Chiapas | FC44 | 0/20 (100%) | 1.9 K | 45.1 B | 1-3 |
| Jalisco | FC22 | 0/20 (100%) | 1.3 K | 48.8 A | 1-3 |

The average incubation period of *CLso* in tomato landraces ranged from 21.5 to 48.8 dpi. The genotypes FC22, FC44, FC40, and FC33 showed a significantly (*P* = 0.0001) greater delay in the first symptoms of *CLso* with an average between 25.6 and 48.8 dpi in comparison with the rest of the genotypes that had an average incubation period between 21.7 and 22.9 dpi “Table 1”. The time of appearance of the first symptoms correlated negatively and significantly with the average values of the symptoms in the studied genotypes (r = -0.814, *P* = 0.004).

Most of the tomato landraces and susceptible controls used in this study did not have a wide variation in symptom levels of *CLso*. Landraces FC22, FC44, FC40, and FC33 showed a range of symptoms of 1-3, 1-3, 3-9, and 3-9, respectively, whereas the rest of the genotypes had a range of symptoms of 7 -9 “Table 1”.

Origin, genotype, incidence of disease: number of resistant plants (NPR) and total tested plants (TTP), average of symptoms severity (Means), time of *CLso* incubation in days (Incubation), and range of studied symptoms in a scale from 1 to 9 (Range) at 60 days post inoculation. Means with the same upper-case letter within columns indicate non-significant differences (*P* ≤ 0.05). NA: not applied.

### Second resistance assay

In this assay, the genetic resistance to *CLso* was analyzed only with genotypes that showed at least one resistant plant during the first trial to corroborate its resistance level under the same conditions of the previous test. Most of the plants inoculated with *B. cockerelli* were susceptible to *CLso* infection. However, landraces FC22 and FC44 were again the most resistant landraces showing significantly greater percent of resistant plants, less symptoms severity, lower *CLso* titers, and longer incubation time, in comparison with the rest of the genotypes “Table 2”. Landraces FC40 and FC33 were fully susceptible in this trial and showed no significant difference with the Pony Express and Reserva susceptible controls for all parameters measured in this test “Table 2”.

**Table 2.**
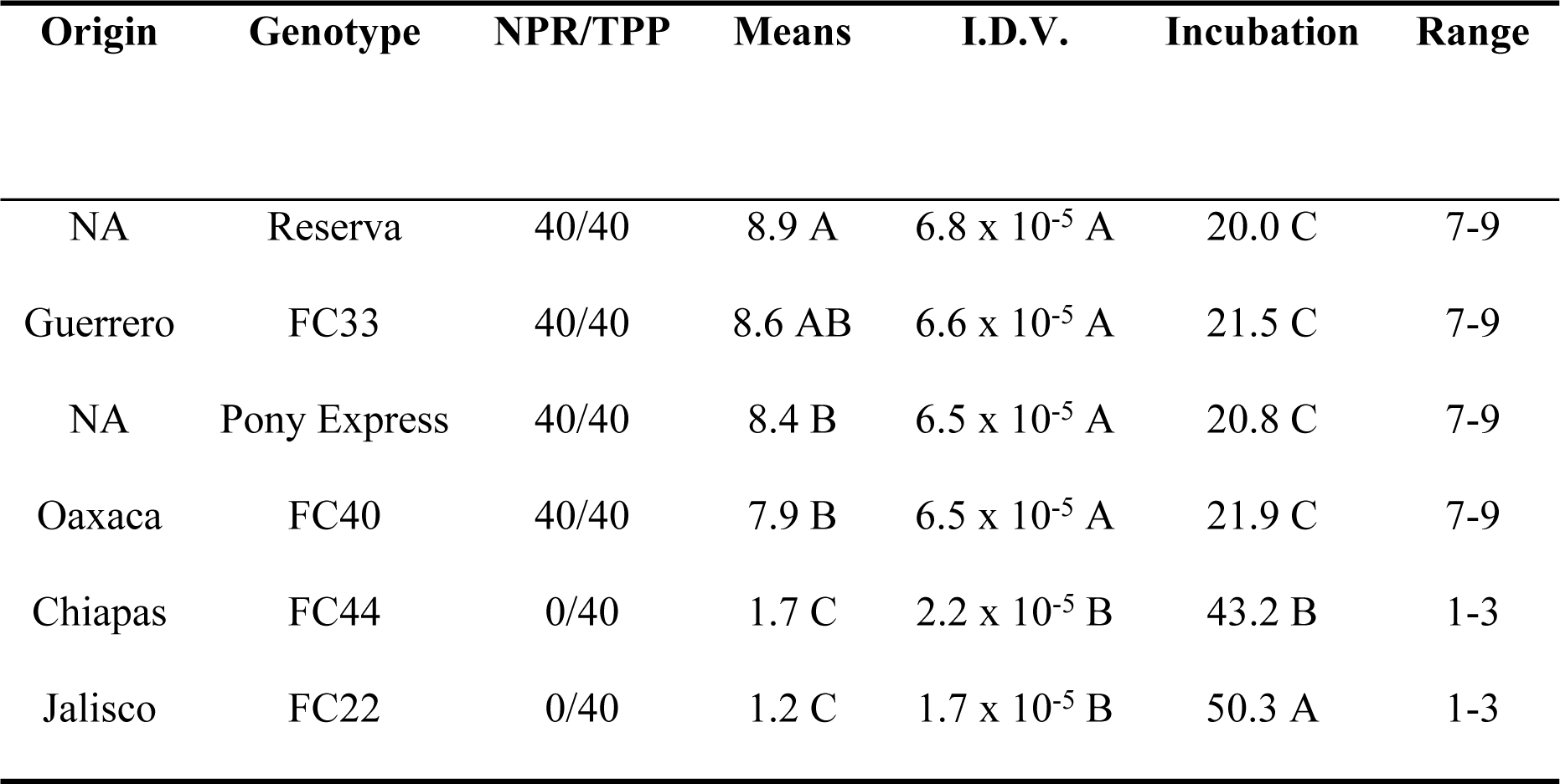
Second resistance assay of landraces FC22, FC44, FC40, FC33, and the cultivars Pony Express and Reserva inoculated with *CLso* through *B. cockerelli*.

In this trial, the average incubation period of *CLso* in tomato landraces ranged from 20.0 to 50.3 dpi. Landraces FC22 and FC44 showed a significantly (*P* = 0.0001) longer incubation period of symptoms in comparison with genotypes FC40, Pony Express, FC33, and Reserva which had the shortest delay in developing the first symptoms of *CLso* “Table 2”. Again, the time of appearance of the first symptoms correlated negatively and significantly with the average values of the symptoms in the studied genotypes (r = -0.734, *P* = 0.007).

Most of the tomato landraces used in this trial did not have a wide variation in *CLso* symptom levels. Landraces FC22 and FC44 showed both a range of symptoms of 1-3, while the rest of the genotypes (FC40, Pony Express, FC33, and Reserva) had on average a range of symptoms of 7 -9 “Table 2”.

The insects identified as *B. cockerelli* used in this study achieved a 100% transmission efficiency in both assays with 20 adults per plant with the “Mexico Sinaloa, *B. cockerelli*” strain of *CLso*. In addition, *CLso* DNA was detected in 100% of all resistant plants analyzed in both assays.

Origin, genotype, incidence of disease: number of resistant plants (NPR) and total tested plants (TTP), average of symptoms severity (Means), relative quantity concentration of *CLso* DNA, where mean values are expressed in integrated density values (I.D.V.), time of *CLso* incubation in days (Incubation), and range of studied symptoms in a scale from 1 to 9 (Range) at 60 days post inoculation. Means with the same upper-case letter within columns indicate non-significant differences (*P* ≤ 0.05). NA: not applied.

## Discussion

*CLso* is one of the most challenging bacterial species to manage in tomato crops because its control is based mainly on chemicals through the use of insecticides against its vector insect *B. cockerelli* and the restriction for the use of these products has been gradually incremented globally. On the other hand, chemical control is partially effective, costly, represents a biohazard, and contribute to the development of resistance in vector populations [15-17]. However, an effective alternative, without bio-risk, and accepted for the integrated management of this bacterium is the development of resistant genotypes [13,16,18,19]. Therefore, to help reduce the *CLso* problematic in tomato in Mexico and other parts of the world, the implementation of breeding programs needs to be considered.

On the other hand, this bacterium is a relatively new pathogen described, attacking tomato plants in New Zealand, Central and North America during the last 10 years [1,3-8]; probably, tomato breeding programs from the seed companies have had delays in launching new cultivars with different levels of resistance to this bacterium due to the lack of resistance sources, genetic studies of this resistant trait, molecular markers, and studies regarding this problematic that would encourage them to invest in it.

Mexico is reported to be one of the potential centers of origin for domesticated *Solanum lycopersicum* species [22-24]. Plant centers of origin often possess the highest genetic diversity and are considered potential sources of resistant genotypes against plant diseases and insect pests [30-32]. Based on that, we hypothesize that landraces from this country must have different level of resistance to *CLso*.

From 46 *S. lycopersicum* landraces used in this study 44 were susceptible, and 2 (FC22 and FC44) showed high levels of resistance to *CLso* according to the low level of symptoms described by [14], the long incubation period of the first symptoms, and the low bacterial titers showed during two consecutively trials in different years “Table 1 and 2”. These results indicate that these landraces are promising sources of resistance to *CLso*. In addition, these genotypes with high levels of resistance can be used individually for the development of resistant tomato cultivars to this bacterium and/or to further improve resistance through the complementation of these sources of resistance to create lines and cultivars with high level and more durable resistance to this important and problematic bacterium. Our results are similar to those reported by [14,33] who found high levels of resistance to *CLso* in different accessions of *S. lycopersicum* from Mexico.

Tomato landraces used in this study had a relative narrow variation in *CLso* resistance levels. Landraces FC22 and FC44, which were resistant in both trials and developed significantly lower levels of symptoms in comparison with the rest, showed a narrow variation in the level of symptoms between them with a range of symptoms from 1 to 3. However, this variation in resistance made it possible to find resistant plants with low indices of symptoms within each landrace and these results were corroborated during the second trial in the search for resistance against this bacterium “Table 1 and 2”. This relative narrow variation in resistance is not in agreement with those reported by [26,27], who found a wide genetic variation in several morphological plant traits of tomato landraces from Mexico. These differences could be attributed to the different traits measured in these studies and to the different tomato backgrounds of the genotypes.

The methodology to discriminate between susceptible and resistant genotypes was optimal because the insects used in this study achieved a 100% transmission efficiency of *CLso* in all plants of both assays with 20 adults per plant and because *CLso* DNA was detected in 100% of all resistant plants analyzed in this study, indicating that the inoculation method was adequate to discriminate between susceptible and resistant plants. This approach can be used in further genetic studies to develop molecular markers and to support tomato breeding programs in the developing of cultivars resistant to *CLso*. These results agree with [14,33,40], who reached 100% efficiency of *CLso* transmission with 20 adults of *B. cockerelli* per plant during the inoculation of tomato and potato plants.

Tomato landraces FC22 and FC44, which were resistant in both trials, are promising sources of resistance to *CLso*. Further studies to analyze the genetic base of this resistant trait must be carried out to design the best genetic model for the introgression of this *CLso*-resistance trait into cultivated tomato backgrounds.

## Materials and Methods

### Plant material

Fruits were collected from around 10 plants of each of the 46 populations of tomato landraces used in this study during the autumn-winter season of 2015 in the states of Sinaloa, Nayarit, Jalisco, Michoacán, Guerrero, Oaxaca, and Chiapas. Pony Express (HM-Clause) and Reserva (Vilmorin) tomato crops were used due to their high susceptibility to *CLso* [14,33]. The seeds were germinated in trays of 200 polystyrene cavities in a germination chamber at 30 °C ± 2 °C.

### Source of inoculum and maintenance of the vector insect

*CLso*-infected *B. cockerelli* insects were collected according to the morphological characteristics reported by [34] with an entomological aspirator from tomato plants (Pony Express cv.) showing classical symptoms of this bacterium under open field conditions. Insects were reared on tomato (Reserva cv.) in growth chambers, in 150 × x 50 × 100-cm insect-proof wooden cages with 60 × 60 mesh organza fabric to avoid contamination of other insect vectors, at temperatures ranging between 26 and 30 °C, at a 16:8 photoperiod (L:D) to increment and conserve the inoculum source and insects during 3 months prior to our experiments. Tomato plants were replaced as needed.

### Identification of *CLso*

For *CLso* identification, DNA samples from four plants of the commercial tomato field infested with *B. cockerelli*, plus four samples from the insects colony (samples of 10 insects each) were tested by PCR using primer pairs Oa2/OI2CF and Oa2/OI2CD that amplify a sequence from the 16S rDNA gene of 1168 bp [3]. DNA from the analyzed insects and plants was extracted following the method of [35]. PCR analysis used for *CLso* detection followed the description by [3].

One positive amplicon from one plant sample of the susceptible control “Pony Express” of all samples analyzed by PCR was excised from the agarose gel and purified through silica columns (EZ-10 Spin Column BS354 DNA Gel Extraction Kit, Bio Basic Inc., Markham, ON, Canada). After obtaining the purified PCR fragment, the sample was sent for sequencing to the National Laboratory of Genomics for Biodiversity (LANGEBIO) of the Center for Research and Advanced Studies (CINVESTAV-IPN), Irapuato, GTO, Mexico. A similarity search with the DNA sequence was performed using the BLAST program, whereby the nucleotide sequences in the study were compared with those in the GenBank databases (NCBI); http://www.ncbi.nlm.nih.gov/BLAST/).

Phylogenetic analyses were performed using the 16S rDNA gene sequences. The sequences from all the isolates were aligned using Clustal W [36]. A neighbor-joining tree was generated based on the alignment obtained with MEGA 7.0 [37]. The confidence of the nodes was tested with 1000 bootstrap replicates.

### Resistance assays

In this study, two *CLso* tests were performed. The first assay was done in March 2016. For the insect inoculation, the methodology used was that reported by [14], which consists of placing a plastic bottle with 20 viruliferous insects at adult stage on individual plants with 2-3 true leaves age of the different genotypes for a 48-h transmission period. After inoculation, an imidacloprid (Confidor®, Bayer Crop Sciences) treatment was applied to eliminate insects. The study was conducted under a completely randomized design with 46 treatments (*S. lycopersicum* landraces) with 20 replicates of one plant per genotype. Five plants of each genotype were not inoculated and were considered as negative control. These control plants were covered individually with large mesh bags for the 48-h inoculation period to prevent any vector contamination into the greenhouse. Plants were maintained under greenhouse conditions as described above and fertilized as required. Two months after inoculation, all the plants were scored. The resistance to *CLso* was assessed based on the level of symptoms severity of the inoculated plants at 60 days post inoculation (dpi) using the scale proposed by [14] with some modifications. Disease severity was rated on a scale of 1 to 9, where: 1 = plant without symptoms; 3 = plant with light yellowing; 5 plant with moderate yellowing, epinasty, filimorfism, rolling and crispy leaves; 7 = plant with strong yellowing, epinasty, filimorfism, rolling and crispy leaves; 9 = stunted plant with all these symptoms expressed strongly or dead plant. Plants that obtained a score equal or less than 3.0 were considered as resistant.

The second assay was performed in March 2017 with four landraces that showed at least one resistant plant from the first trial to corroborate their level of resistance. The same methodology of inoculation with 20 insects, number of plants per genotype, experimental design and scale to determine resistant plants applied in the first trial was used in this assay.

All plants from both trials were analyzed for the presence of *CLso* DNA by PCR to discard escape plants. Both trials were done under greenhouse conditions with temperatures fluctuating between 20 (night) and 31 ± 2 °C (day), and a 12:12 h photoperiod.

### Quantification of *CLso* DNA

*CLso* DNA was quantified by a densitometry method (IS-100 Digital Image System Alpha Innotech Corporation, San Leandro, CA, USA) and expressed as integrated density values (I. D.V.). This quantification was carried out on the four landraces that showed at least one resistant plant to *CLso* in the first test and on two susceptible cultivars (Pony Express and Reserva). Three plants of each infected genotype were assessed.

### Statistical analysis

Data obtained from the assessments of genetic resistance to *CLso* from both assays were subjected to non-parametric variance analysis with the Kruskal–Wallis test and Dunn median to determine the significance among genotypes (*P* = 0.05). All statistical analyses were performed with the SAS software [38].

## Acknowledgments

Authors thank FitoCiencia, for the support provided to this research.

## Author Contributions

Conceptualization: José Antonio Garzón-Tiznado, Luciano Castro-Espinoza

Formal analysis: Carlos Alfonso López-Orona, Sixto-Velarde-Félix

Methodology: José Antonio Garzón-Tiznado, Jesús Enrique Retes-Manjarrez

Resources: Jesús Enrique Retes-Manjarrez

Supervision: Marely Graciela Figueroa-Pérez, Jesús Enrique Retes-Manjarrez

Writing ± original draft: Jesús Enrique Retes-Manjarrez

Writing ± review & editing: José Antonio Garzón-Tiznado, Jesús Enrique Retes-Manjarrez

